# Transcription imparts architecture, function, and logic to enhancer units

**DOI:** 10.1101/818849

**Authors:** Nathaniel D Tippens, Jin Liang, King Y Leung, Abdullah Ozer, James G Booth, John T Lis, Haiyuan Yu

**Author notes:** contributed equally (Nathaniel D Tippens, Jin Liang). correspondence should be addressed to (JL) and (HY).

## Abstract

Distal enhancers remain one of the least understood regulatory elements with pivotal roles in development and disease. We used massively parallel reporter assays to perform functional comparisons of two leading enhancer models and find that gene-distal transcription start sites (TSSs) are robust predictors of enhancer activity with higher resolution and specificity than histone modifications. We show that active enhancer units are precisely delineated by active TSSs, validate that these boundaries are sufficient to capture enhancer function, and confirm that core promoter sequences are required for this activity. Finally, we assay pairs of adjacent units and find that their cumulative activity is best predicted by the strongest unit within the pair. Synthetic fusions of enhancer units demonstrate that adjacency imposes winner-takes-all logic, revealing a simple design for a maximum-activity filter of enhancer unit outputs. Together, our results define fundamental enhancer units and a principle of non-cooperativity between adjacent units.

## Introduction

Since their identification in viral and mammalian genomes, enhancers have been defined primarily by their function: the ability to activate promoters independently of their distance and orientation^1–3^. More basic questions about the nature of enhancer elements remain difficult to answer: what are the genomic features of active enhancers? How large are they? Classical examples such as the α- and β-globin locus control regions (LCRs) offer some clues: these LCRs are predominantly driven by 400-900 bp DNase hypersensitive sites (DHSs) harboring transcription factor (TF) binding and extensive non-coding transcription^4,5^. These features were also observed from all enhancers identified from a recent CRISPR-Cas9 screen of the *MYC* locus^6^. Histone modifications such as H3K27ac^7^ and H3K4me1^8^ have been proposed to mark enhancers, although such predictors lack systematic comparison^9–11^. Similarly, genome annotation tools such as ChromHMM^12^ have been developed using histone modifications to generate enhancer predictions averaging 600 bp in size.

The finding that transcription from distal enhancers is widespread and corresponds with activation^13,14^ led to numerous hypotheses about roles and functions of non-coding “enhancer” RNAs (eRNAs). Many long non-coding RNAs (lncRNAs) were thought to facilitate gene-regulatory functions, but systematic introduction of premature polyadenylation signals demonstrated that most lncRNAs are dispensable; instead, recruitment of transcription and splicing complexes drives their gene-regulatory function^15,16^. Recently, a “molecular stirring” model has been proposed wherein transcription increases molecular motion that drives enhancer-promoter interactions^17^. Similarly, we have proposed that RNA Polymerase II’s (RNAPII) affinity for common co-factors or even itself could facilitate enhancer-promoter interactions^18,19^. This model is supported by reports that the C-terminal domain (CTD) of RNAPII specifies active promoter localization through its affinity for other CTDs^20^, as well as the low-complexity domain of Cyclin T1^21^. If correct, these models suggest that transcription may be required for distal enhancer function, challenging the commonplace methodology of using DNase hypersensitive sites (DHSs) and histone marks to identify enhancers. More fundamentally, functional enhancer transcription would imply structure within enhancer sequences because transcription requires well-positioned core promoter sequences for assembly of the pre-initiation complex^22^.

Numerous high-throughput sequencing methods identify enhancers using either plasmid or integrated reporter constructs and are collectively known as massively parallel reporter assays (MPRAs). While these assays offer unprecedented throughput for surveying genome function, their technical biases and limitations are a focus of ongoing research and optimization^23–25^. For example, most published MPRAs have been limited to short synthetic sequences (50-150 bp), despite the precise size of genomic enhancers remaining unknown^11^. The development of Self-Transcribing Active Regulatory Region sequencing (STARR-seq) circumvented this limitation with a simple cloning strategy to quantify genomic fragments as large as 1,500 bp by placing them into the 3’ untranslated region (3’UTR) of a reporter gene^2^. After transfecting cells with the reporter library, enhancers will drive their own RNA expression. Each candidate’s enhancer activity is then defined as the ratio of mRNA to plasmid DNA, as quantified by Illumina sequencing.

In this study, we perform systematic functional comparisons of commonly used histone marks versus transcription initiation patterns that are frequently observed at enhancers. We find that transcription is found at virtually all active distal enhancers and validate a basic unit model for enhancers defined by their TSSs. Finally, we establish approaches for quantifying unit cooperativity and uncover a position-encoded mechanism by which stronger enhancers overshadow adjacent enhancer units.

## Results

Seven MYC enhancers that were recently identified by CRISPR-Cas9 interference exhibit many conventional features of active enhancer architecture^6^. For example, *MYC* enhancer 2 (segment A) is a DNase I hypersensitive site (DHS) and contains elevated levels of H3K27ac and H3K4me3 (Figure 1a). It also contains a single divergent TSS pair. To test features critical for enhancer function, we sub-cloned (segment C) from the larger A region previously verified by luciferase assays, as well as flanking sequences (segments B & D) for comparison. Notably, segment C harbored virtually all observed distal enhancer activity in luciferase assays (Figure 1b). A nearby site with similar DNase hypersensitivity and histone modifications that does not exhibit divergent transcription (segment E) did not show significant enhancer activity. This example illustrates how divergent transcription may help localize active enhancer boundaries with high resolution, and avoid ambiguities derived from lower-resolution DNase and chromatin immunoprecipitation (ChIP) profiles.

**Figure 1.**
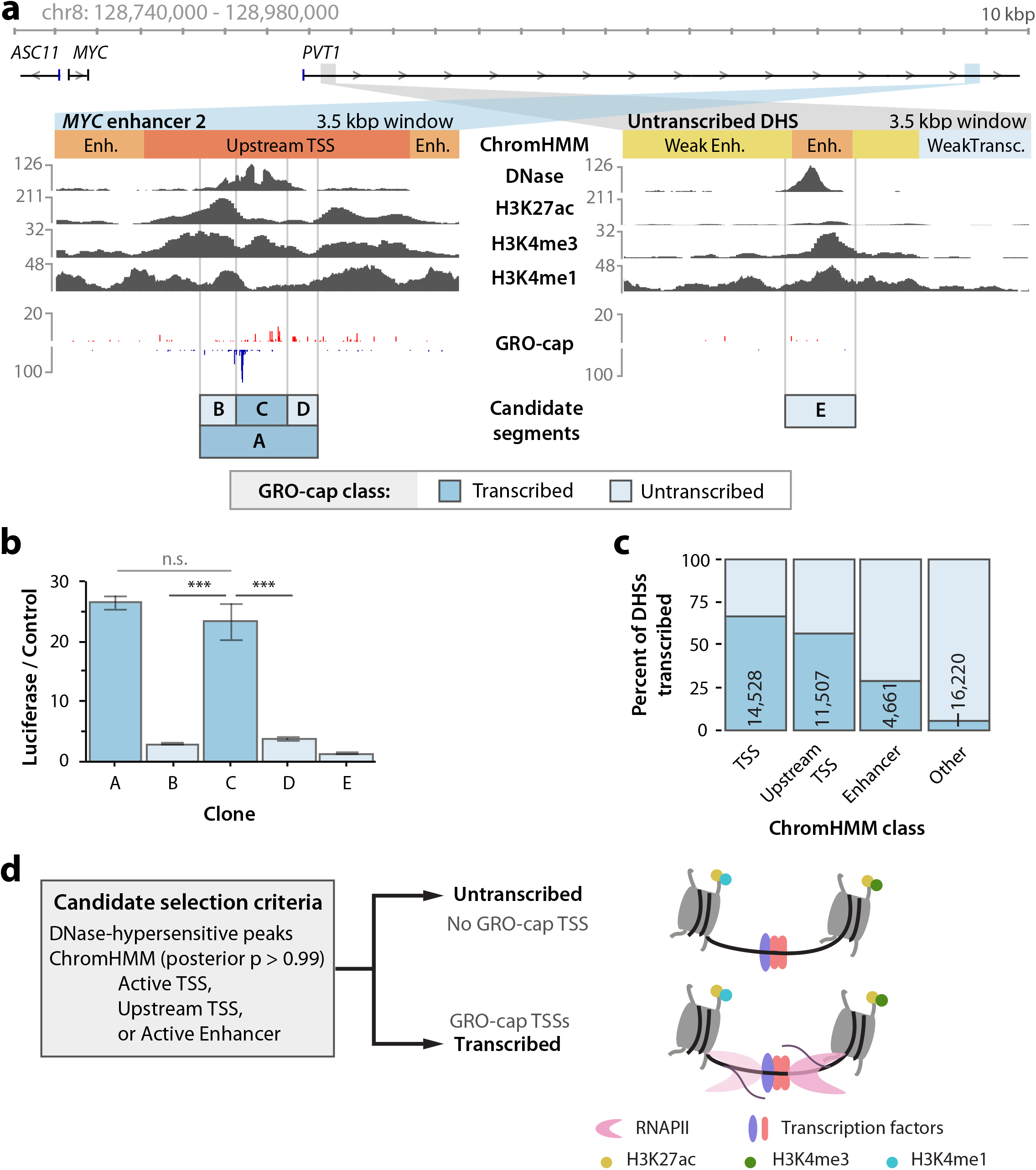
Divergent transcription identifies enhancer boundaries in high resolution. **a.** Genomic data tracks of two candidate regulatory elements in the MYC locus. Raw read counts are shown for each track, and the “Candidate elements” track indicates cloning boundaries used for luciferase assays of tested sequences. **b.** Renilla-normalized luciferase reporter activity for the regions indicated in **a**. Error bars indicate standard error of the mean. **c.** The percent of DHSs within each indicated ChromHMM class that are untranscribed (no GRO-cap TSS call) vs transcribed (containing GRO-cap TSS call). Number of transcribed DHS are indicated. **d.** A schematic of candidate element selection using DNase, ChromHMM, and GRO-cap data. Molecular model compares DHS sharing many features, with or without RNAPII transcription. n.s. = not significant, p > 0.1; *** = p < 0.0005; student’s t-test.

To generalize these results, we systematically sampled a larger set of potential enhancers in K562 cells. This set was chosen to include DHSs from combinations of active ChromHMM classes^12^, and transcription initiation classes defined by Global Run-On Cap data^19,26^ (GRO-cap; see Methods). Notably, most DHSs do not contain a GRO-cap TSS (86%). However, DHSs from the Active Enhancer, Active TSS, and Upstream TSS ChromHMM classes are enriched for one or more GRO-cap TSSs (Figure 1c). We compare enhancer activity of transcribed and untranscribed DHSs from only high-confidence examples of these ChromHMM classes (posterior p > 0.99; Figure 1d). Selected candidates ranged from 180-300 bp in size (Figure S1a).

### Divergent transcription marks active enhancer elements

In order to test hundreds of candidate enhancer sequences, we adapted STARR-seq for use with sequence-verified elements as large as ~2 kbp, which we call element-STARR-seq (eSTARR-seq; Figure 2a). We clone every candidate sequence in both forward and reverse orientations within the 3’UTR of the reporter gene to distinguish sequences that may regulate mRNA stability. We added unique molecular identifiers (UMIs) to the reverse transcription primer for removal of PCR duplicates, and tagmentation before library amplification to circumvent the length limitations and minimize biases of Illumina sequencing (Figure 2a; see Methods). As in other MPRAs, enhancer activity is quantified as the ratio of mRNA to transfected DNA (after de-duplication with UMIs). eSTARR-seq improves agreement with luciferase data compared with conventional STARR-seq (Figure S1b), likely because UMIs increase the dynamic range, and is highly reproducible from true biological replicates (Figure 2b). Finally, we measure the relationship between fragment size and reporter activity using negative controls consisting of human open reading frames (ORFs), which are unlikely to destabilize mRNA or harbor distal enhancer activity (Figure S1c). In conclusion, eSTARR-seq enables robust quantification of enhancer activity while minimizing PCR, size, and orientation biases.

**Figure 2.**
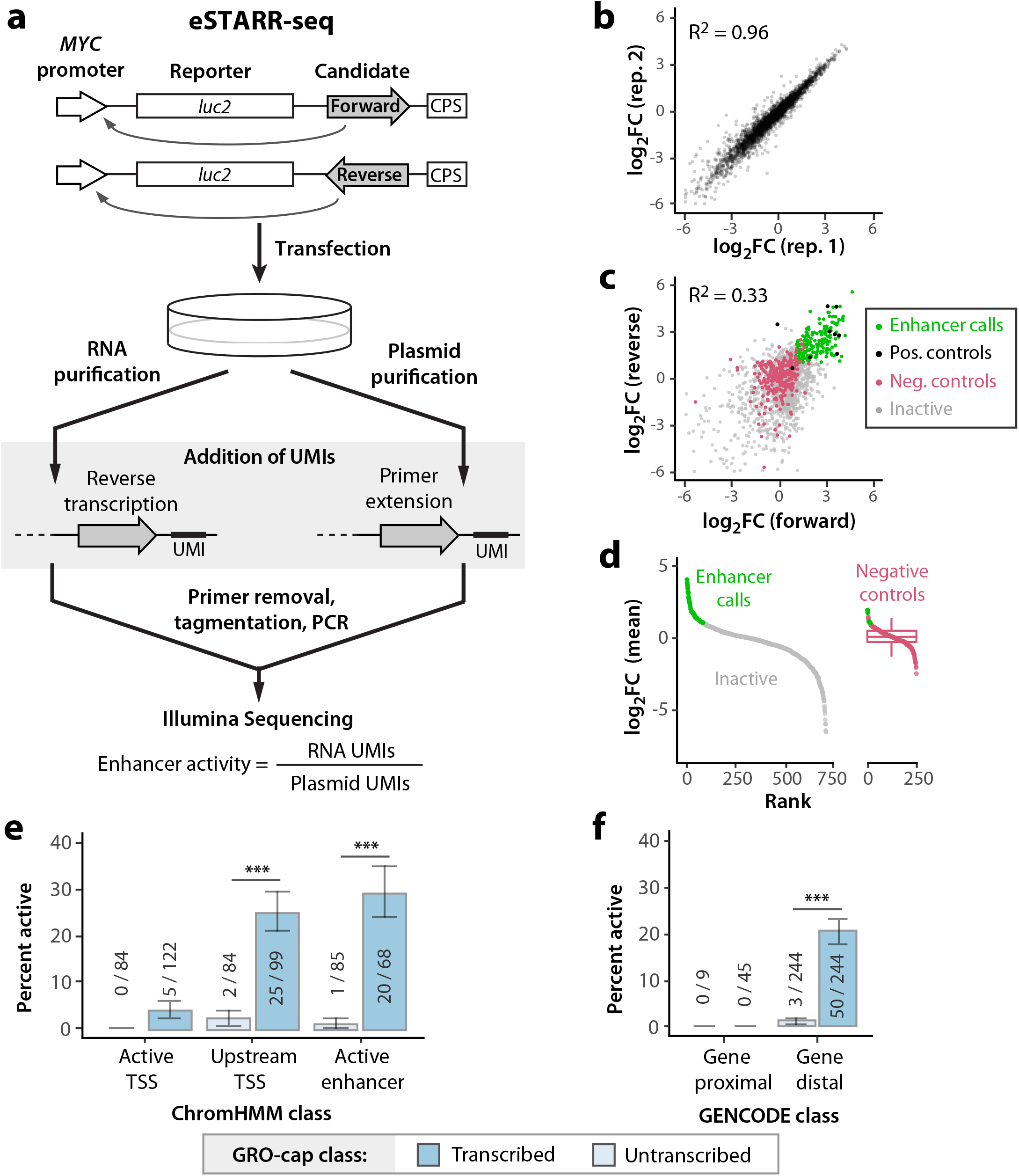
Transcription robustly predicts active eSTARR-seq enhancers. **a.** Outline of element-STARR-seq (eSTARR-seq). Each candidate is cloned into the 3’UTR of a luc2 reporter gene in both forward and reverse orientations. After transfection into K562 cells, total RNA and plasmids are purified separately. Addition of unique molecular identifiers (UMIs) occurs during reverse transcription for RNA, or primer extension for plasmids. After sequencing, enhancer activity is estimated by the ratio of RNA to plasmid UMIs. **b.** eSTARR-seq is highly reproducible between biological replicates. **c.** Comparison of estimated activity from forward vs reverse cloning orientations. Data points are shown as log2 fold-change vs negative controls, averaged from three replicates. Positive controls are known MYC or viral enhancers (black). Negative controls are human open reading frames (ORFs, magenta). Elements with significantly elevated activity in both orientations are called enhancers (green). Remaining candidates are called inactive (gray). **d.** Summary of enhancer calls made in **c** after averaging forward and reverse activities. Empirical false-discovery rate is 2.4% (6/243 negative controls misidentified as enhancers). **e-f.** Within each ChromHMM (**e**) or GENCODE (**f**) class, the percent of active enhancers identified by eSTARR-seq is indicated. “Gene proximal” is defined as within 500 bp of a GENCODE protein-coding transcript 5’ end. Error bars indicate standard error calculated for a sample of proportions.*** = p < 0.0005; N-1 Chi-square test.

Enhancer activity is known to be orientation-independent^1,3^, whereas mRNA stability is affected by strand-specific RNA sequences. Thus, we required candidates to exhibit significantly higher reporter activity than controls in both forward and reverse cloning orientations to be classified as an enhancer (Figure 2c; see Methods). Only 2.6% (6/243) of negative controls met these criteria, confirming very few false-positive enhancer calls (Figure 2d).

Comparing transcribed and untranscribed DHS revealed that most eSTARR-seq activity was found in transcribed DHSs from the Upstream TSS and Active Enhancer ChromHMM classes (Figure 2e). Within these two classes, 25-30% of transcribed candidates exhibited significant enhancer activity (compared with ≤ 2% for untranscribed candidates). Importantly, GRO-cap provides similar predictive performance without ChromHMM after using a 500 bp distance cut-off from GENCODE annotations to distinguish gene promoters from distal enhancers (Figure 2f). These results significantly extend recent reports^10,27–29^ by demonstrating that virtually no active enhancers are untranscribed when using the most sensitive nascent TSS methods such as GRO-cap, and strongly suggests a possible functional role for transcription from active enhancers.

### Transcription delineates regulatory sequence architecture

Given the striking co-occurrence of transcription initiation and active enhancer elements, we revisited the model that promoters and enhancers share a universal architecture^13,30^ (Figure 3a). Classic studies defined minimal “core promoter” sequences that coordinate assembly of the pre-initiation complex^22^; here, we define core promoters as beginning 32 bp upstream of the TSS (the location of TFIID binding to the TATA box motif when present^22^) and ending at the RNAPII pause site (≤60 bp beyond the TSS^19^). Two distinct core promoters are found up to 240 bp apart (corresponding to ~300 bp between TSSs) and may help position the −1 and +1 nucleosomes^31^. By contrast, the “upstream region” contains regulatory TF motifs that may activate one or both core promoters.

**Figure 3.**
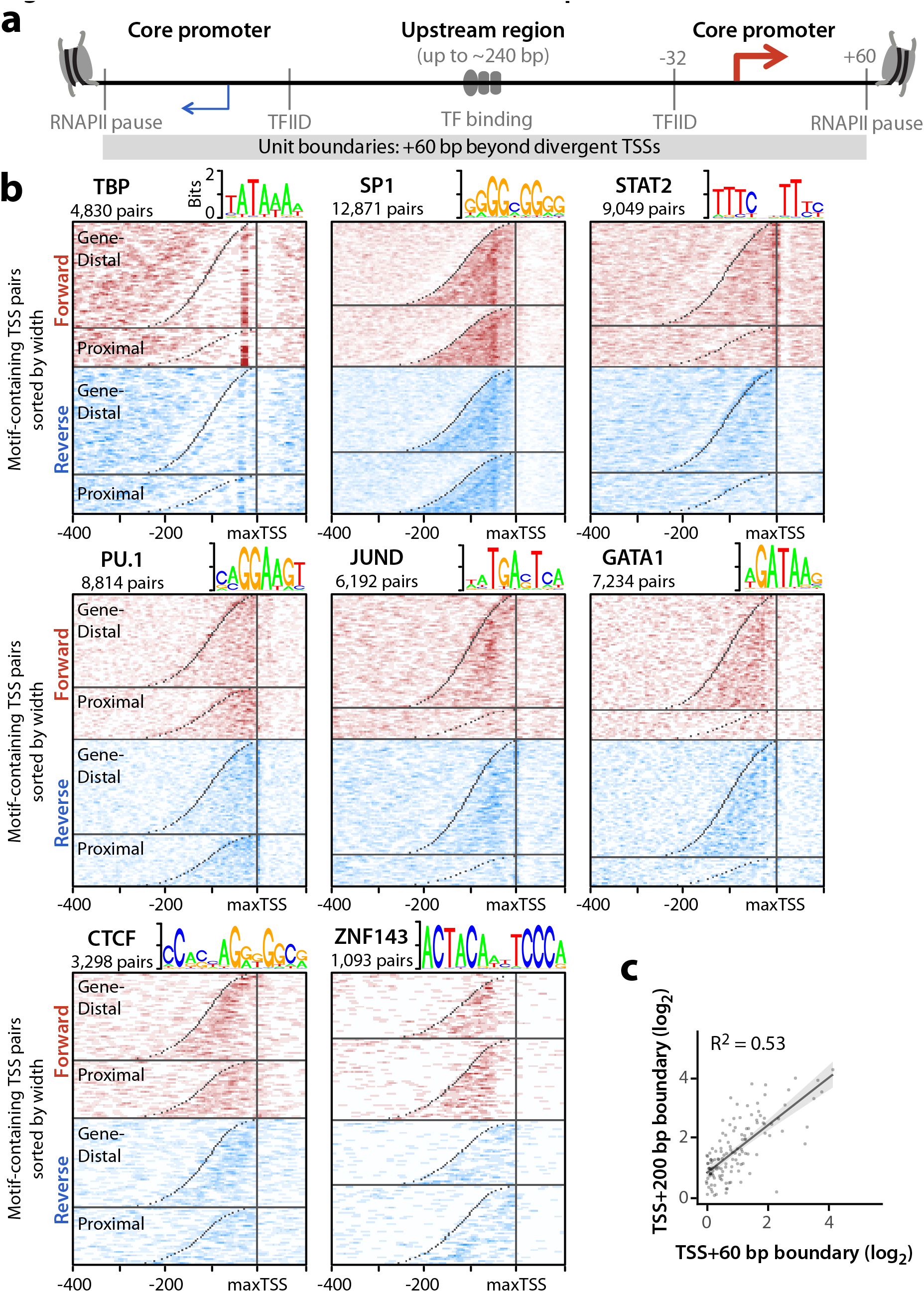
Enhancer unit boundaries reveal sequence architecture. **a.** Illustration of a unified model for regulatory sequence architecture of promoters and enhancers. Core promoter motifs (TBP, SP1, STAT2) surround an upstream region containing TF motifs. We define core promoters as the region from transcription factor II D (TFIID) binding 32 bp upstream of each TSS, to the RNAPII pause sites at +60 bp from each TSS. **b.** Divergent TSS pairs were sorted by width and aligned to the max TSS. TSS pairs were also divided by GENCODE class (Gene-distal vs Gene-proximal). Heatmaps indicate TF motif densities from pairs containing at least one motif. Motifs are shown in both forward (red) and reverse (blue) orientations relative to the max TSS. TSS positions are marked in gray. **c.** Comparison of enhancer activities for the same set of elements using TSS+60 bp and TSS+200 bp cloning boundaries. Line of best fit is shown with 95% confidence interval shaded gray.

To illustrate similarities in sequence architecture at both promoters and enhancers, we plotted motif densities relative to the stronger TSS for both distal and gene proximal TSS pairs (Figure 3b). Interestingly, some motifs are well-aligned to TSSs, especially those known to recruit and position TFIID. Similar to the well-known TATA-box bound by TBP, SP1^22^ (max motif density at −53 bp), and STAT2^32^ show striking TSS alignment and are known to recruit TFIID. Systematic classification of core promoter sequences is particularly important since <10% of human TSSs contain a TATA box, and a recent report demonstrated that core promoters respond differently to co-activators and distal enhancers^22,33,34^. However, most motifs appear dispersed throughout the “upstream region” between divergent TSSs, as illustrated by PU.1, JUND and GATA1. By contrast, CTCF and ZNF143 motifs are found near the weaker TSS. Notably, CTCF and ZNF143 have been implicated in facilitating distal loop interactions, reinforcing the idea that similar motif alignments identify similar regulatory mechanisms. Whereas ChIP-seq analyses can only reveal central and core promoter binding TFs^13^, sequence motif analyses reveal more nuanced spatial preferences within these elements^35^.

We re-tested a subset of elements after adding 140 bp of sequence context on each side to test whether core promoter boundaries are sufficient to capture enhancer activity (TSS+60 bp vs TSS+200 bp). Importantly, adding sequence context affected enhancer activity less than testing identical fragments in differing orientations (Figure 3c R^2^=0.53 compared with Figure 2c R^2^=0.33). This indicates enhancer activity appears to be fully captured with sequences extending 60 bp beyond divergent TSSs, thus providing a basic unit definition of enhancers. In summary, we validate our boundary definition of individual enhancer units and reveal motif alignments that might help decipher regulatory function^34–36^.

### Enhancer units require core promoters for activity

Next, we sought to determine whether all components of the divergent TSS model (Figure 3a) are necessary to drive distal enhancer activity. If transcription is spurious and unimportant to enhancer function, core promoter sequences should be dispensable. To answer this question, we re-analyzed the “High-resolution Dissection of Regulatory Activity” (HiDRA) dataset^37^, which uses the STARR-seq assay on Analysis of Transposase-Accessible Chromatin (ATAC-seq) fragments. This impressively comprehensive dataset quantifies enhancer activity from 100-600 bp fragments enriched within DHSs, thus dissecting potential enhancer elements genome-wide. Given our observations of pronounced orientation effects in STARR-seq assays (Figure 2c), we re-analyzed the HiDRA dataset to remove this bias wherever possible. Unfortunately, most HiDRA fragments (87%) do not share significant overlap with a fragment tested in the opposite orientation (Figure 4a). We assessed orientation bias across all 763,373 fragment pairs tested in both orientations with ≥90% overlap and found very little agreement across orientations (Figure 4b; HiDRA R^2^ = 0.07). Interestingly, HiDRA fragments that contain a DHS exhibit less orientation bias (Figure S2a; R^2^ = 0.38), closely matching our eSTARR-seq results (R^2^ = 0.33; Figure 2c).

**Figure 4.**
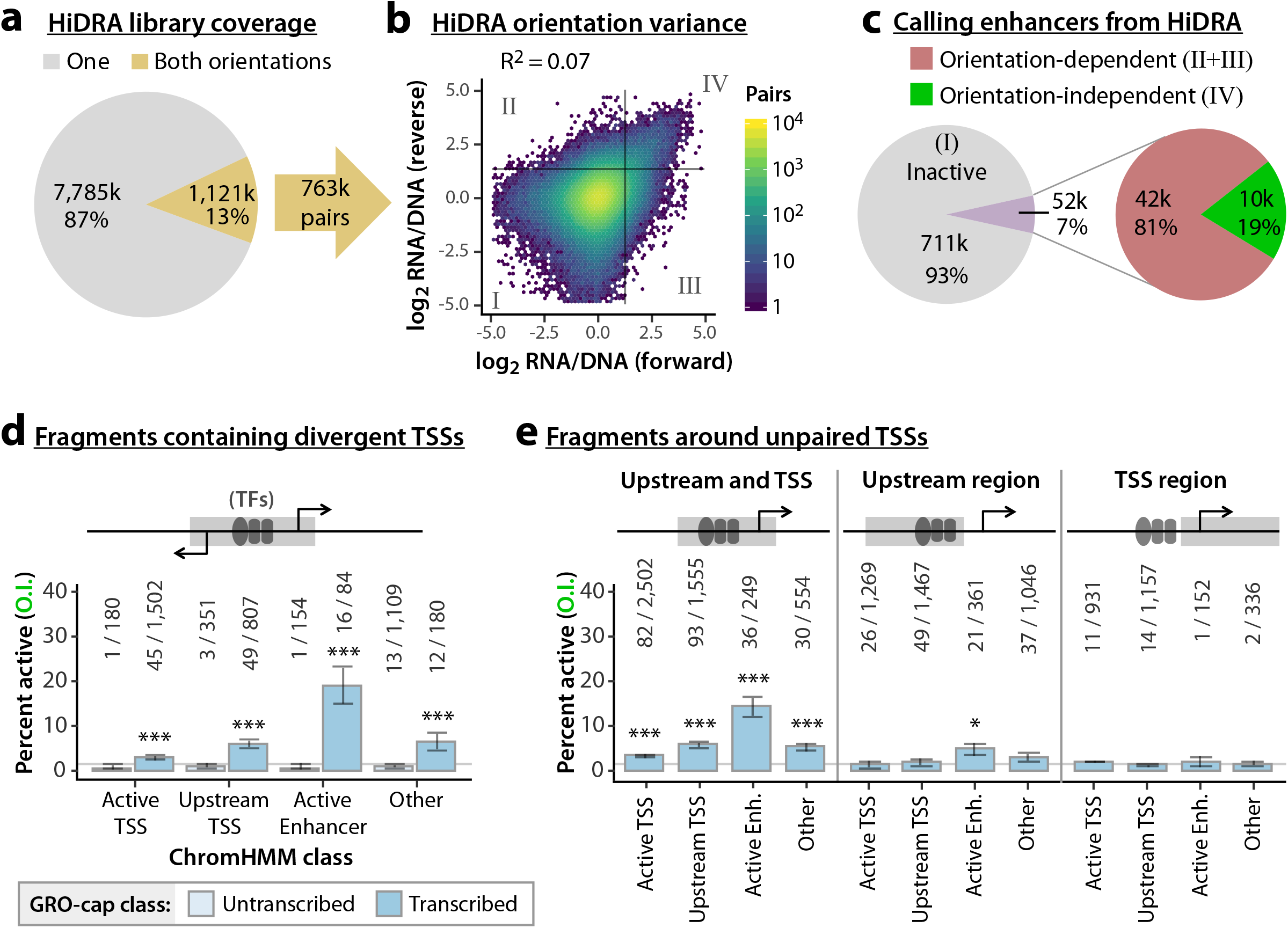
Upstream and TSS regions are both necessary for enhancer activity. **a.** Pie chart indicating the fraction of HiDRA fragments tested in one (gray) or both (gold) orientations. Some fragments have pairings with more than one fragment in the opposing orientation, providing 763,000 distinct pairs. **b.** Comparison of HiDRA enhancer activities from opposing orientations of fragment pairs. Color indicates the number of pairs. Gray lines denote approximate statistical cut-off for active enhancers. Quadrants II and III denote orientation-dependent “enhancer” fragment pairs; quadrant IV fragments are active in both orientations. **c.** Pie chart indicating the percent of HiDRA fragment pairs classified as inactive, orientation-dependent, and orientation-independent. **d-e.** Bar charts indicating the percentage of orientation-independent enhancer calls from HiDRA fragments sample from DHSs within the indicated ChromHMM classes. **d,** fragments are further classified as untranscribed or transcribed (contains divergent GRO-cap TSSs). **e,** fragments are sampled from different areas around unpaired GRO-cap TSSs (see cartoon and Methods). Raw fragment counts are shown above each bar. Gray line marks the average percent activity of all fragments. Error bars indicate standard error calculated for a sample of proportions. * = p < 0.05; ** = p < 0.005; *** = p < 0.0005; N-1 Chi-square test.

Importantly, accounting for orientation bias has substantial impact on enhancer identification. While 93% of HiDRA fragment pairs appear inactive (Figure 4b, Quadrant I), the 7% of fragment pairs with elevated RNA/DNA signal (Quadrants II-IV) are dominated by orientation bias (Quadrants II-III): only 19% of these fragment pairs exhibit elevated activity in both cloning orientations (Quadrant IV, Figure 4c). This is true even when only considering fragments that span a DHS, with 71.2% of enhancers exhibiting orientation-dependence (N=580/827 enhancer fragment pairs; Figure S2a). Interestingly, most transcribed DHSs showed enrichment for orientation-dependent activity (Figure S2b). When using stringent orientation-independent enhancer calls, HiDRA identifies only 0.22% of tested fragments as enhancers, although we predict this should be improved by selection of larger fragments to increase capture of whole elements.

HiDRA fragments containing enhancer units defined by divergent TSSs were most enriched in the Active Enhancer ChromHMM category (Figure 4d), confirming our observations in K562 cells (Figure 2d). To determine if one or both core promoter sequences are necessary for enhancer activity, we computed the fraction of HiDRA enhancers around unpaired GRO-cap TSS. At these sites, the upstream and TSS regions can be easily separated from each other (Figure 4e). Strikingly, we observed little enrichment for orientation-independent enhancers from upstream or TSS regions alone, while activity is strongly enriched within fragments containing both the TSS and upstream regions (Figure 4e). These results demonstrate that core promoter sequences within TSS regions are necessary for distal enhancer activity, and strongly suggest a functional role for RNAPII recruitment to enhancers. Our findings are reminiscent of recent dissections of promoter activity^38^ and provide strong support for similar sequence architectures at promoters and enhancers^13,30^, although they each exhibit clearly distinct functionalities (Figure 2e).

### Proximity-encoded logic regulates neighboring enhancer elements

Many gene-distal TSSs are found in dense regulatory clusters that have complex histone modification patterns^19^, implying widespread clustering of basic enhancer units. To explore how individual enhancer units (subunits) might cooperate within these clusters, we fit a model to predict the enhancer activity of a cluster from its subunits’ activities (Figure 5a). 100 clusters and associated subunits were successfully cloned so that their enhancer activity could be quantified independently within the same experiment. 45% of clusters showed significant enhancer activity compared with negative controls (Figure S3a), and predominantly contained a single active sub-element (Figure S3b).

**Figure 5.**
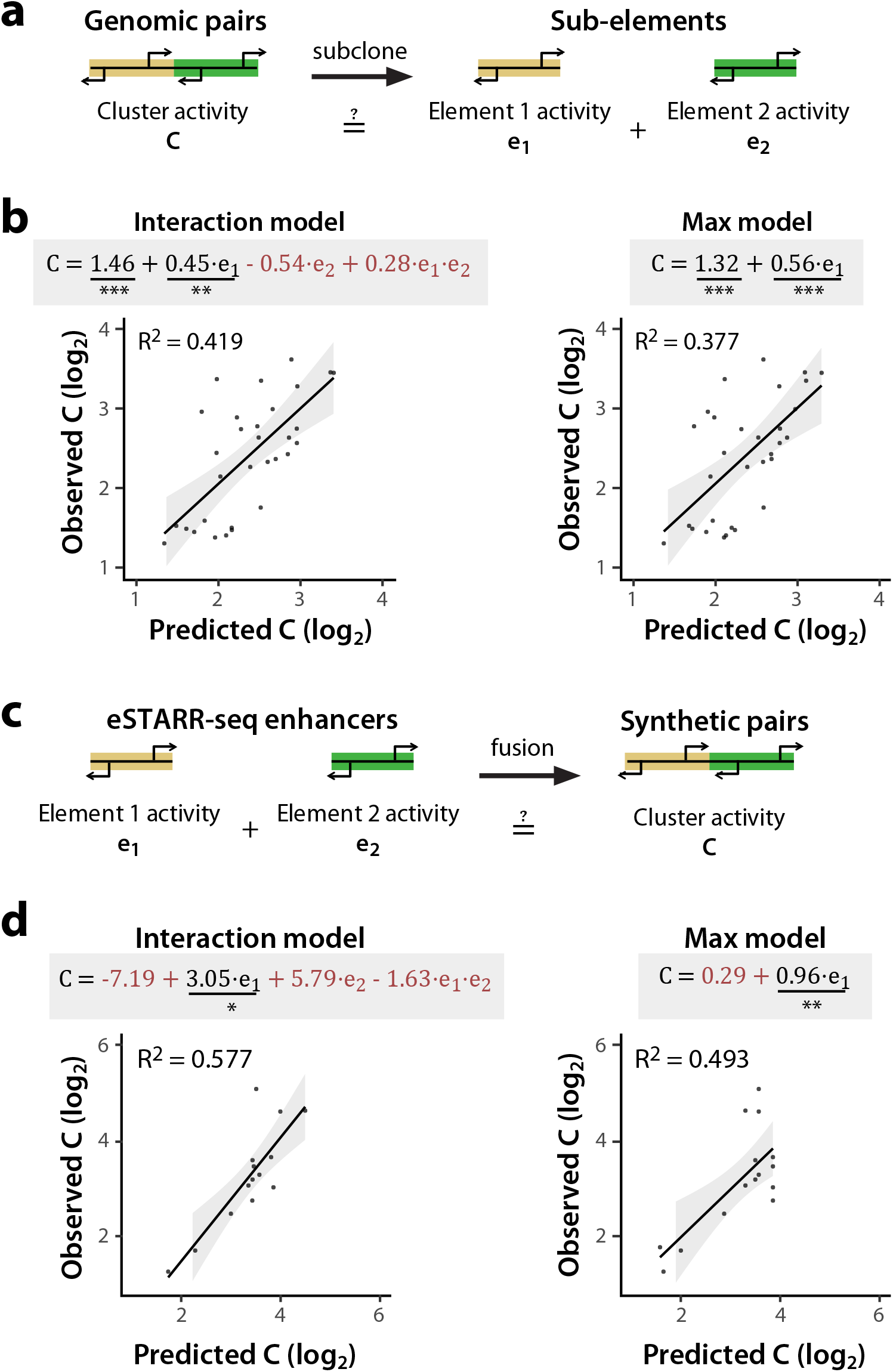
Adjacent enhancers are non-cooperative. **a.** Dissection of genomic TSS clusters into individual sub-elements to quantify enhancer cooperativity. **b.** Two linear models were fit to eSTARR-seq measurements of full clusters (C) and individual enhancers within the cluster (e1 and e2). The interaction model includes both individual enhancers and an interaction term, while the max model only considers the stronger sub-element (chosen to be e1). Fitted equations are shown with significant covariates underlined and non-significant covariates colored red (ANOVA). Shaded area denotes 95% confidence interval for the line of best fit. **c.** Schematic illustrating fusion of active enhancer sequences into synthetic enhancer pairs. **d.** Fitting of same linear models in b to enhancer activities of individual elements and their synthetic fusion (as shown in **c**). * = p < 0.05; ** = p < 0.005; *** = p < 0.0005.

We fit a linear model to predict cluster activities (Interaction model, Figure 5b) from the observed subunits’ activities (e_1_ and e_2_) and an interaction term (e_1_×e_2_). Strikingly, this analysis revealed significant covariance between cluster activity and the subunit with higher activity (p=0.0004), but not the subunits with lower activity. Indeed, including only the subunit with higher activity (Max model) explains 37.7% of the observed variance (Figure 5b), which was not significantly less than the Interaction model (p = 0.14). This suggests that clusters could be predominantly driven by a single subunit, or that neighboring enhancers target different promoter classes.

To directly quantify enhancer unit cooperativity, we generated synthetic pairs made by randomly fusing eSTARR-seq active enhancer units (Figure 5c). This targeted approach removes the possibility that the cluster’s subunits are communicating with different promoters, since we can select pairs where both enhancer subunits were already found to drive the promoter of our reporter construct. We developed a pooled strand-overlap extension PCR strategy to fuse units into random pairs linked with a constant 25 bp sequence. This method generated 138 fusions, 15 of which were pairs of active enhancer units (Figure S4a). Individual units were re-tested in the same pool as the fused sequences, and their eSTARR-seq activities agreed well with previous measurements (Figure S4b). Surprisingly, the interaction model including both subunits still did not find statistically significant predictive power from the weaker subunit and failed to outperform the Max model (p = 0.15), demonstrating that proximity to a stronger enhancer effectively abolishes weaker enhancers’ activity. The max model explains 49.3% of the variance among active enhancer pairs, and 36.3% of the variance among all enhancer-containing pairs (N=68; Figure S4c). As expected, the Max model does not perform well for pairs lacking any enhancer activity, explaining only 16.9% of the variance (N=70; Figure S4d). These results demonstrate that immediate proximity of enhancer units in DNA allows only the strongest enhancer to function, and therefore encodes a max-activity filter likely regulating dense enhancer clusters genome-wide.

## Discussion

Although transcription and histone modifications are closely correlated^8,11,13^, we find that histone marks are lower resolution and less specific for enhancer activity^9,10^ than transcription initiation patterns provided by GRO-cap^13,26^. We further demonstrate that TSSs are useful anchors in revealing motif positioning within enhancers^35^ and enable dissection of regulatory clusters into individual subunits.

Previous analyses of conserved enhancers across species found widespread TF motif rearrangements that did not impact function, leading to a “flexible” sequence model for enhancers that was only evaluated with promoter-proximal MPRAs^39,40^. Using data from a distal enhancer reporter assay, we find that enhancer activity requires at least one core promoter in addition to specific TF combinations in the flexible upstream region, suggesting a functional role for RNAPII recruitment at enhancers. Likewise, recent analyses of population variants affecting gene-distal GRO-cap TSSs suggest that core promoter mutations in distal enhancers can disrupt enhancer function^28^. The requirement for core promoters at enhancers is particularly intriguing given reports that core promoters confer specificity for enhancers and co-activators^22,33,34^; this suggests enhancers could conceivably target promoters through recruiting similar core promoter machinery. Additionally, RNAPII pausing at enhancers^10^ may facilitate distal interactions through the CTD’s affinity for other CTDs^20^, resulting in coordinated pause release at promoters and associated enhancers by P-TEFb kinase^41^. Further analysis of regulatory sequence architectures at promoters and enhancers may expand the lexicon for non-coding elements beyond individual TF motifs and clarify enhancer-promoter interaction specificities and mechanisms.

Consistent with previous studies^2,24,29^, we find few gene promoters with distal enhancer activity, despite striking similarities in their chromatin architecture. This highlights lingering questions about the distinguishing features of these two regulatory elements. In general, promoters and enhancers have been reported to differ in GC content and TF recruitment preferences, but such rules lack specificity^30^. Core promoter sequence features might help distinguish enhancers from promoters, particularly if RNAPII itself reads a regulatory code during pausing and/or early elongation. For example, RNAPII pausing is sequence dependent^19,42^, and is substantially longer-lived at promoters than enhancers^10^. Stable RNAPII pausing at promoters may provide time to recruit distal regulatory complexes by co-localization with the unstable RNAPII pausing seen at enhancers. Finally, burst size is encoded within core promoter sequences^43^. Promoters may favor selection for larger burst sizes, whereas enhancers maximize burst frequency to drive distal gene activation^44^.

Recently, enhancer clusters have been dissected *in vivo* resulting in different models of their cooperativity^45–47^. Statistical re-analysis of these data demonstrated that both reports are consistent with multiplicative generalized linear models^48^, although statistical power was greatly constrained by sample size. While these studies assessed cooperativity over significant distances (2-50 kbp), we assayed dozens of adjacent enhancer pairs (≤600 bp apart) and fit a single multiplicative (or log-additive) linear model to explain their cumulative activity. Our dataset surveys a much larger number of clusters and indicates non-cooperativity between adjacent elements, revealing a simple design for a max-activity filter of enhancer outputs. Indeed, a recent report of alternative TSS selection within distal enhancers during differentiation underscores broad implications of the max-activity filter^49^. This regulatory mechanism provides evolution a versatile tool for cellular decisions through winner-takes-all logic and may be easily adaptable for genetic engineering applications in agriculture and medicine.

## Author contributions

N.D.T, J.L., A.O., J.T.L., and H.Y. conceived of the project and designed the enhancer comparison screen. N.D.T. conceived of dissecting enhancer cooperativity and mechanisms. J.L. performed most cloning, primer design, and all eSTARR- and Clone-seq assays. N.D.T. optimized the pooled stitch PCR protocol and prepared enhancer fusions with guidance from A.O., H.Y., and J.T.L.. N.D.T. and K.Y.L. performed analysis with feedback from J.G.B., J.L., A.O., J.T.L, and H.Y.. N.D.T. wrote the manuscript with feedback from all authors.

## Competing Interests

None.

## Data Availability

Processed GRO-cap data was obtained from Gene Expression Omnibus expression GSE60456. Raw sequencing files for the HiDRA study were obtained from SRA accession SRP118092. eSTARR-seq data is being made available through the ENCODE data portal (accession pending). All candidate regulatory element clones generated in this study and used for eSTARR-seq and luciferase assays are available upon request. Please address requests to Haiyuan Yu.

## Code Availability

All analysis scripts are available on Github (https://github.com/hyulab/eSTARR).

## Methods

### Candidate element selection, definition, and primer design

To systematically compare transcribed and untranscribed candidates within each ChromHMM class, we focused on high-confidence Active TSS, Upstream TSS, and Active Enhancer predictions (posterior p > 0.99). This set of regions was then filtered by requiring overlap with ENCODE DHS peaks from K562 cells (<u>E123-DNase.macs2.narrowPeak.gz</u>). Finally, ChromHMM regions were classified as either transcribed or untranscribed by overlapping with GRO-cap divergent peaks (from supplementary files of reference^13^). ~300 Untranscribed ChromHMM regions were selected for cloning using DHS peak width as boundaries. Similarly, ~600 Transcribed ChromHMM regions were selected for cloning using TSS+60 bp boundaries, where the TSS position was determined as the max GRO-cap signal within the peak. Primers were allowed to be no more than 10 bp from the desired boundaries. See Figure S1A for element sizes within each class.

### eSTARR data analysis

Cutadapt was used to identify attB1 sequences within each read. Next, a custom python script was used to extract element sequences and remove PCR duplicates (identical PCR barcode + first 15 bp of element). Processed reads were then aligned to candidate elements with bowtie2 (--end-to-end-a). A custom R script was used to extract alignments within 3 bp of the expected cloning boundaries, ensure complete removal of PCR duplicates, and generate orientation-specific read counts for each candidate, provided in Supplementary Tables 1-2. All analysis scripts are available on Github (https://github.com/hyulab/eSTARR).

To identify elements with significant enhancer activity, raw read counts were processed using *voom* from the R Bioconductor limma package. RNA and DNA counts were treated as distinct experimental conditions within each replicate. Active enhancers were defined as having significantly elevated ratio of RNA to DNA counts with FDR-adjusted p < 0.1 in both cloning orientations. Additionally, we required log_2_ fold-change ≥ 1 in both cloning orientations to ensure significantly higher activity than negative controls (Figure 2c).

### HiDRA data analysis

Raw sequencing files were obtained from SRA (accession SRP118092) and aligned to the hg19 genome as described^37^ (bowtie2 -p 6, -q and --phred33). BAM files were merged within replicates using samtools, then processed with a custom R script to remove multi-mappers (mapq < 30) and apply size selection (100-600 bp). Differential RNA vs DNA read counts were detected using *voom* from the R bioconductor limma package. To minimize size bias, voom was applied separately to fragments from 100-150 bp, 150-200 bp, etc. After applying voom, we only considered fragments with ≥5 DNA counts (summed from all replicates) to minimize artifacts of low-coverage sites. Alignments with mutual overlap >= 90% and mapping to opposite strands were considered as a “forward” and “reverse” alignment pair. We required FDR-adjusted p < 0.1 in both forward and reverse cloning orientations to call active enhancer fragments. HiDRA enhancer fragments were then analyzed relative to published GM12878 GRO-cap peaks^13^. GRO-cap peaks were collapsed to the single most-used transcription start nucleotide with a custom R script. All analysis scripts are available on Github (https://github.com/ndt26/eSTARR).

For dissection of unpaired GRO-cap TSSs, “Upstream and TSS” fragments were defined as containing at least 200 bp upstream and 30 bp downstream of a GRO-cap TSS. “Upstream region” fragments were taken from between 330 and 35 bp upstream of a GRO-cap TSS. “Core promoter region” fragments were defined to contain at least 40 bp upstream and 190 bp downstream of a GRO-cap TSS.

### Motif density analysis

K562 and GM12878 GRO-cap divergent pairs and processed GRO-cap data were obtained from published work^13^. Peaks were refined to a single nucleotide according to the maximum GRO-cap signal within each TSS. Divergent pairs were required to be less than 300 bp apart for visualization. Genomic sequences from −400 to +100 bp of the max TSS of each divergent pair were scanned for motifs using RTFBSDB with default match settings^50^. This scan produces in an Nx500 count matrix, where N is the number of sites scanned, and 500 bp is the window size. Each entry in the matrix is 0 (motif absent) or 1 (motif present). After removing divergent pairs without any matching motifs, loci were sorted by distance between their divergent TSSs and whether they were proximal (within 500 bp) or distal to a GENCODE gene annotation start coordinate. Finally, neighboring rows in the count matrix were averaged into 100 groups to compute motif density at each position for each strand, and normalized to the maximum density observed in the matrix. This matrix was plotted at 4 bp resolution for simplicity; most motifs are 8-12 bp. All analysis scripts are available on Github (https://github.com/ndt26/eSTARR). All motif density profiles shown in Figure 3 are from K562 GRO-cap TSSs, except for STAT2, which was derived from GM12878 GRO-cap TSSs.

### eSTARR-seq assay vector

The eSTARR-seq assay vectors were generated by modifying the original STARR-seq vector^2^. To engineer the pDEST-hSTARR-luc-Pmyc vector, the Synthetic Core Promoter (SCP) in the STARR-seq vector was replaced with the *MYC* promoter^6^ and the truncated sgGFP was replaced with a luciferase reporter gene (*luc2*). Additionally, the two cloning sites and the DNA fragment between them in the STARR-seq vector were replaced with an attR1-attR2 Gateway cassette. To engineer the pDEST-hSTARR-luc-Pmyc-ccw vector, the attR1-attR2 Gateway cassette in pDEST-hSTARR-luc-Pmyc vector was removed and then re-cloned back to its original position in the reverse orientation. Additionally, we generated a pDEST-hSTARR-luc vector that is almost identical to the pDEST-hSTARR-luc-Pmyc vector except that a SCP1 promoter^2^ was used instead of the *MYC* promoter.

### TRE cloning and input plasmid library preparation

The primers for cloning TREs were designed in batch with a webtool^51^ and synthesized by Eurofins. Each primer contained a 5′-overhang, attB1′ for the forward primers and attB2’ for the reverse primers. Human gDNA was used as template for the PCR reactions. The amplicons were cloned into pDONR223 vector via Gateway BP reactions. The resulted single-colony derived TRE entry clones were verified by Illumina sequencing as previously described^51^.

All verified TRE entry clones were propagated in LB medium supplemented with spectinomycin. The culture was then pooled together for plasmid extraction with E.Z.N.A. Plasmid Midi Kit (Omega Bio-tek, D6904). The TREs were cloned into eSTARR-seq assay vector via *en masse* Gateway LR reactions to generate the input plasmid library. The input library was propagated in LB medium supplemented with ampicillin and the plasmids were extracted with the E. Z. N. A. Endo-Free Plasmid DNA Maxi Kit (Omega Bio-tek, D6926).

### Cell culture

The K562 cells (CCL-243) were purchased from American Type Culture Collection (ATCC). The cells were maintained in the culture medium composed of the Iscove’s Modified Dulbecco’s Medium (ATCC, 30-2005) supplemented with 10% fetal bovine serum (ATCC, 30-2020) at 37°C with 5% CO_2_. Cells used for different biological replicates were cultured separately.

### eSTARR-seq library preparation

The input library plasmids were electroporated into the K562 cells with Cell Line Nucleofector Kit V (Lonza, VCA-1003). For each electroporation, one million cells were mixed with 20 μg plasmids and 100 μL supplemented Nucleofector Solution V and electroporated with a Nucleofector II device (Lonza) using Program T-016. The electroporated K562 cells were recovered in 2 mL culture medium at 37°C with 5% CO_2_ until harvest.

The electroporated K562 cells were harvested after six hours of recovery. Total RNAs were extracted from the cells with TRIzol Reagent (ThermoFisher Scientific, 15596026) according to the manufacture’s instruction. Reverse transcription was performed with the total RNAs as the template using SuperScript III reverse transcriptase (ThermoFisher Scientific, 18080044). The electroporated plasmids were extracted from the cells as previously described^52^. The 1^st^ primer extension was performed with the extracted plasmids as the template. In parallel, another primer extension reaction was carried out with the input plasmid library used for transfection as the template. Reactions were treated with exonuclease I to remove excess single-stranded primer, followed by purification on a MinElute purification column (QIAGEN, 28004).

The 2^nd^ primer extension was performed with the products of both the reverse transcription and the 1^st^ primer extension as the templates. In the library preparation for fusion TREs, a low-cycle PCR was performed with the products of the 2^nd^ primer extension as templates to add the Illumina sequencing adaptors and the indexing barcodes, followed by the acquisition of 240 bp + 360 bp reads on a Miseq Illumina sequencer. In all the other library preparations, the products of the 2^nd^ primer extension went through a low-cycle pre-tagmentation PCR amplification before being tagmented with Tn5 transposomes^53^. Another round of low-cycle post-tagmentation PCR was performed to add the sequencing adaptors and the indexing barcodes, followed by the acquisition of 1 × 75 bp reads on a Nextseq 500 Illumina sequencer.

### Dual luciferase assay

The selected TREs were individually cloned into eSTARR-seq assay vectors via LR reactions and the resulting library of plasmids was extracted with the E.Z.N.A. Endo Free Plasmid Mini Kit II (Omega Bio-tek, D6950). The plasmids were electroporated into K562 cells with Ingenio Electroporation Kit (Mirus, MIR 50115). For each electroporation, 0.5 million cells were mixed with 1-2 μg plasmids and 50 μL Ingenio Electroporation Solution and electroporated with a Nucleofector II device using Program T-016. The pGL4.75 vector (Promega, E6931) was co-electroporated (10 ng/electroporation) as the internal control. The electroporated K562 cells were recovered in 2 mL culture medium at 37°C with 5% CO_2_ until harvest.

The electroporated cells were harvested after 24 hours of recovery for dual luciferase assay. The assay was carried out with Dual-Glo Luciferase Assay System (Promega, E2920) according to the manufacturer’s instruction. An Infinite M1000 Microplate Reader (Tecan, 30034301) was used to quantify the luminescent signals. Cells electroporated with only pGL4.75 vector or with only pDEST-hSTARR-luc-Pmyc vector were used as the background controls for firefly or *Renilla* luciferase activities, respectively.

### Pooled strand overlap extension (SOE) PCR

Using a multichannel pipette, PCR reactions were prepared by pairing forward and reverse oligos appropriately (e.g. A pairs with B, and C pairs with D). 50 μL PCR reactions were carried out using Phusion DNA polymerase for 28 cycles and annealing at 58°C. Amplicons were double purified using Ampure XP beads according to the manufacturer’s protocol and eluted into 40 μl of ddH_2_O. Each amplicon was quantified in a 96-well plate using the QuBIT dsDNA Broad Range reagents and a flourometric plate reader. A pooled annealing and extension reaction was set up as follows:

| Volume | Component |
| --- | --- |
| 10 μL | 5× HF Buffer |
| 10 μL | 5 M Betaine |
| 1 μL | 12.5 mM dNTP Mix |
| 0.5 μL | Phusion DNA Polymerase |
| 10 nM | Pooled PCR products (Forward linker) |
| 10 nM | Pooled PCR products (Reverse linker) |
| to 50 μL | ddH<sub>2</sub>O |

Denaturation was performed at 95°C for 3 min. Annealing was performed by rapid cooling to 50°C for 3 min. Extension was performed at 72°C for 5 min. The reaction was then cooled to 4°C for 5 min.

A final PCR reaction was performed to specifically amplify stitched products. The SOE-PCR reaction mix from the previous step was used directly without any purification:

| Volume | Component |
| --- | --- |
| 20 μL | 5× HF Buffer |
| 20 μL | 5 M Betaine |
| 2 μL | 10 mM dNTP Mix |
| 1 μL | Phusion DNA Polymerase |
| 5 μL | SOE-PCR Reaction Sample |
| 250 nM | Forward primer |
| 250 nM | Reverse primer |
| to 100 μl | ddH<sub>2</sub>O |

Amplification was performed for 8 cycles to minimize bias. Denaturation was 95°C for 3 min, annealing was 65°C for 2 min, and extension was 72°C for 1 min. SOE-PCR amplicons were then size-selected from a nondenaturing 6% polyacrylamide gel.

## Notes

https://github.com/ndt26/eSTARR

