## Supplementary Figures for "Transcription imparts architecture, function, and logic to enhancer units"

### Supplemental Figure 1. Design and validation of eSTARR-seq and selected candidates.

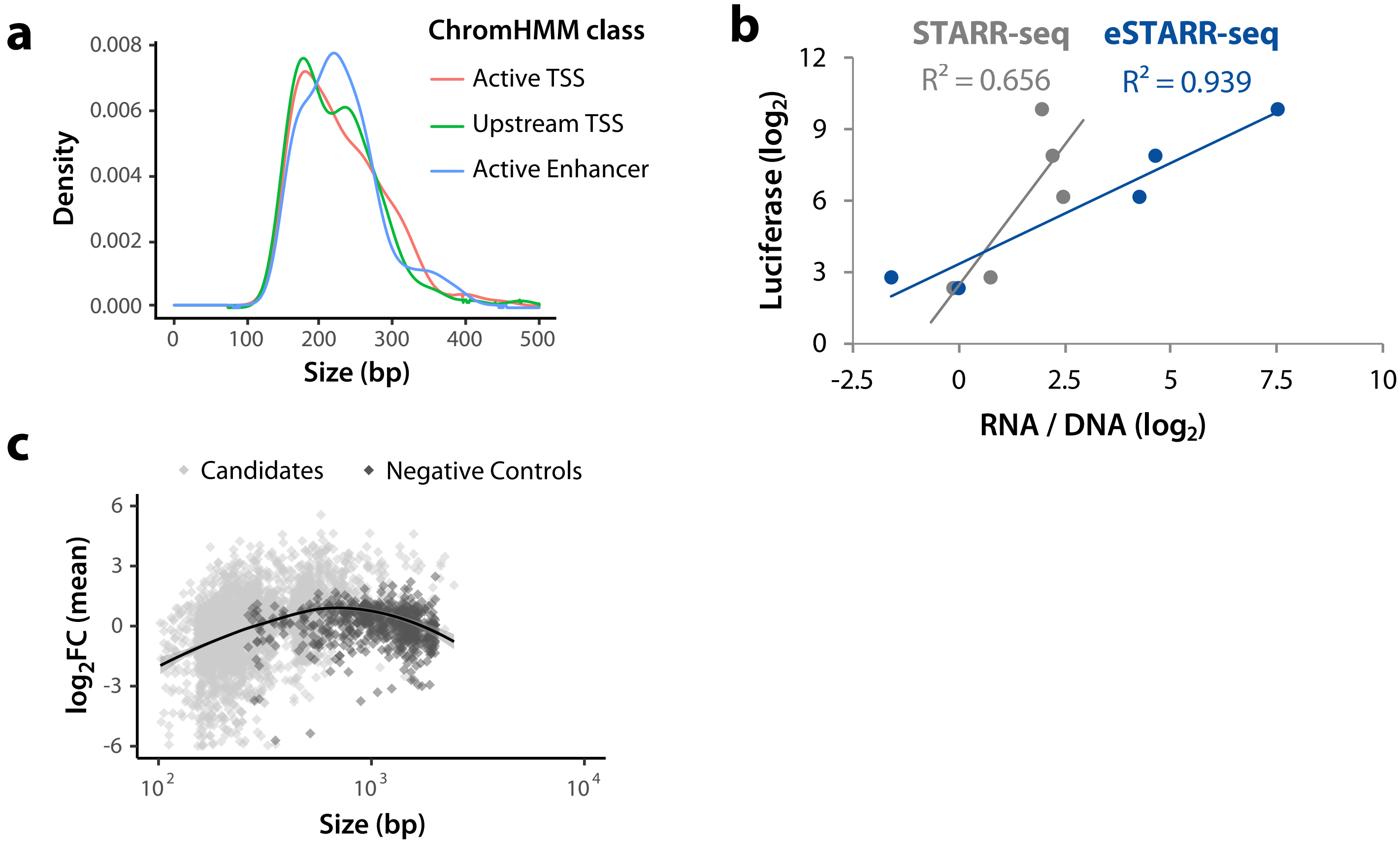


**a.** Size distribution of candidates is shown by ChromHMM class.
**b.** Correlation between conventional luciferase assays and eSTARR-seq in HeLa cells demonstrates greater dynamic range than conventional STARR-seq. Luciferase and STARR-seq activity are from (Arnold et al., 2013).
**c.** Log2 fold-change of elements' RNA/DNA activity vs negative controls is shown relative to each elements' size. Line indicates a fitted loess curve estimate of size bias for eSTARR-seq enhancer activity.

#### Supplemental Figure 2. Orientation dependence in the HiDRA dataset.

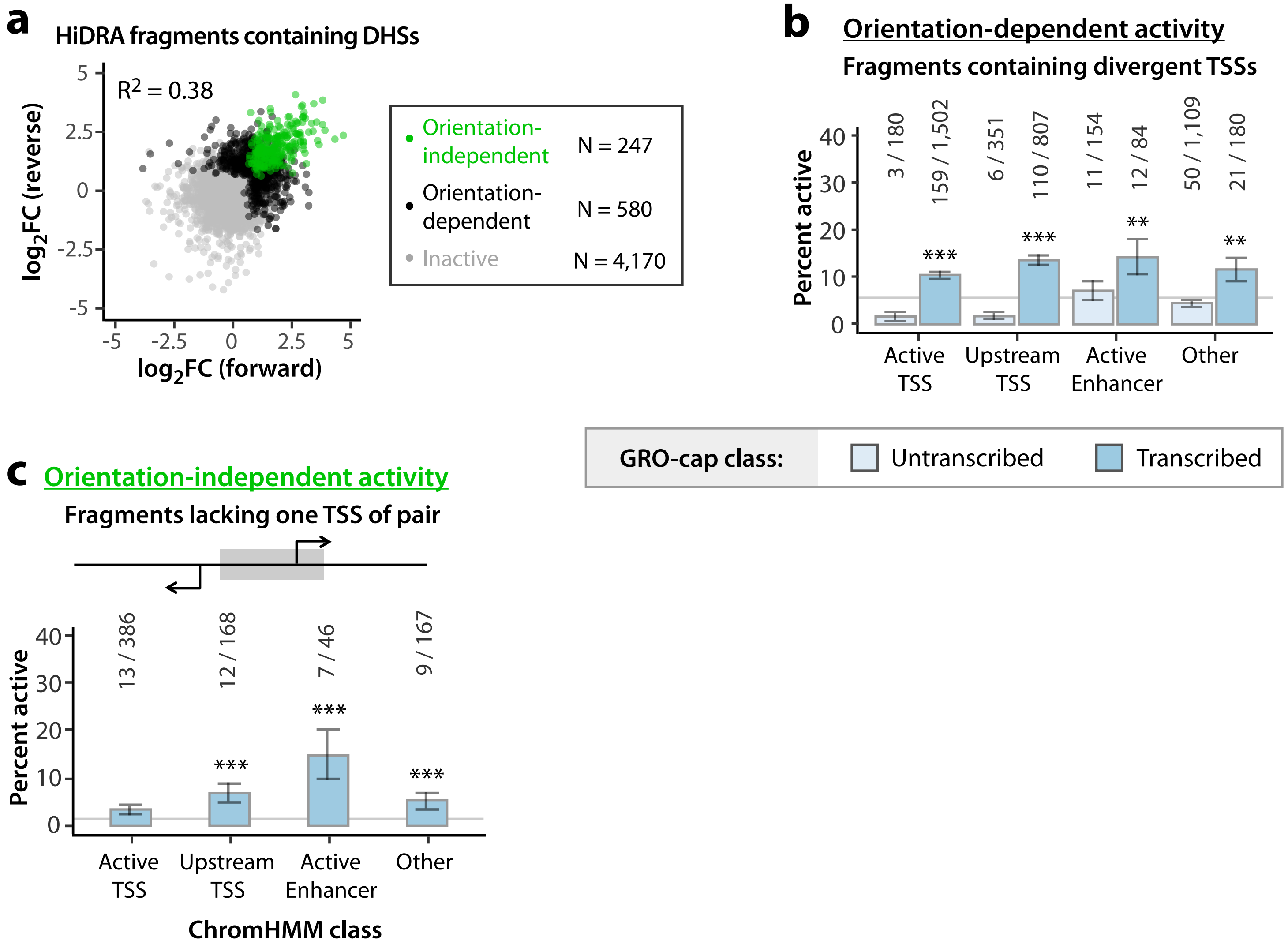


**a.** Comparison of forward vs reverse cloning orientation for HiDRA fragments overlapping GM12878 DHS peaks. Data points are shown as log2 fold-change of RNA vs DNA read counts. Elements with significantly elevated activity in both orientations are called orientation-independent enhancers (green). Elements with significantly elevated activity in one orientation are called orientation-dependent (black). Remaining fragments are called inactive (gray).
**b-c.** Percent of orientation-dependent (**b**) or -independent (**c**) fragments within each GRO-cap and ChromHMM class. Raw fragment counts are shown above each bar. Gray line marks the percent activity of all fragments judged by the same criteria. All error bars indicate standard error calculated for a sample of proportions.
* = $p < 0.05$; ** = $p < 0.005$; *** = $p < 0.0005$; N-1 Chi-square test.

#### Supplemental Figure 3. Functional dissection of genomic TSS clusters.

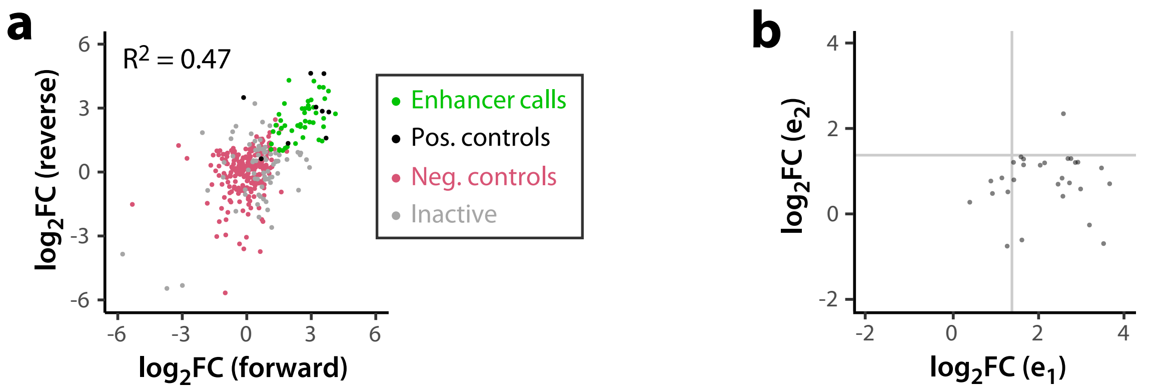


**a.** Comparison of forward vs reverse cloning orientation for all tested TSS clusters. Data points are shown as log2 fold-change vs negative controls (magenta), averaged from three replicates. Positive controls (black) are known *MYC* or viral enhancers. Clusters with significantly elevated activity in both orientations are called enhancers (green). All other tested sequences are called inactive (gray).
**b.** Comparison of sub-element activities within active enhancer clusters. The stronger sub-element is always chosen to be e1, and the weaker sub-element is e2. Gray lines indicate approximate significance cut-offs.

#### Supplemental Figure 4. Design and validation of synthetic unit pairs.

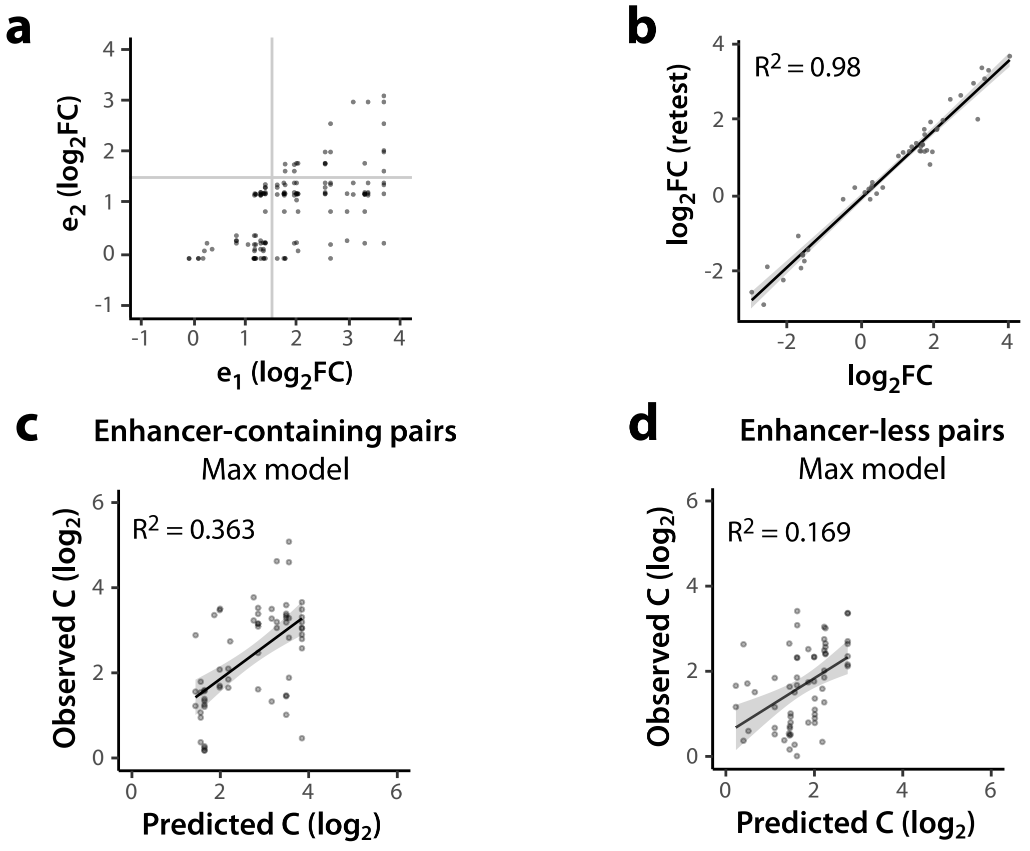


**a.** Comparison of sub-element activities within synthetic enhancer clusters. The stronger sub-element is always chosen to be e1, and the weaker sub-element is e2. Gray lines indicate approximate significance cut-offs.
**b.** Correlation between individual unit eSTARR-seq activities tested previously and re-tested as controls in the synthetic fusion screen.
**c.** Agreement between predicted and observed cluster activities (C) for enhancer-containing synthetic pairs.
**d.** Agreement between predicted and observed cluster activities (C) for enhancer-less synthetic pairs.
